# Critical Role of Histone Tail Entropy in Nucleosome Unwinding

**DOI:** 10.1101/494328

**Authors:** Thomas Parsons, Bin Zhang

## Abstract

The nucleosome is the fundamental packaging unit for the genome. It must both remain tightly wound to ensure genome stability while simultaneously being flexible enough to keep the DNA molecule accessible for genome function. The set of physicochemical interactions responsible for the delicate balance between these naturally opposed processes have not been determined due to challenges in resolving partially unwound nucleosome configurations at atomic resolution. Using a near atomistic protein-DNA model and advanced sampling techniques, we calculate the free energy cost of nucleosome DNA unwinding. Our simulations identify a large energetic barrier that decouples the outer and inner DNA unwinding into two separate processes, occurring on different timescales. This dynamical decoupling allows the exposure of outer DNA at a modest cost to ensure accessibility while keeping the inner DNA and the histone core intact to maintain stability. We also reveal that this energetic barrier arises from a delayed loss of contacts between disordered histone tails and the DNA, and is, surprisingly, largely offset by an entropic contribution from these tails. Our study uncovers the balance of energetic and entropic contributions that dictate nucleosome stability, suggesting that tilting this balance may be a previously-unknown mechanism for regulating genome function.

## Introduction

One important task that every cell faces is to organize its genome into a confined nucleus while keeping the genetic information accessible (1, 2). Eukaryotes overcome this challenge by packaging the genome into chromatin, a dynamic structure that can adjust its compactness depending on the functional state of the underlying DNA (3–5). Chromatin, in turn, is composed of nucleosomes - 147 base pairs of DNA wrapped 1.7 times around a histone octamer in a lefthanded superhelix (6, 7), with the wraps of the DNA commonly termed as the inner and outer layers. As the fundamental organizational unit of the genome, the nucleosome plays a crucial role in all DNA templated processes, including transcription, replication, repair, etc. Using extensive computer simulations, we quantitatively investigate the nucleosome stability and provide high-resolution structural characterizations of the nucleosome disassembly pathway.

Formation of nucleosomes helps to fit the genome inside the nucleus, but also occludes the binding of protein molecules to the DNA. Therefore, the first step in virtually all molecular processes that control the genome function involves the unwinding of nucleosomal DNA, a dynamical transition that has been the focus of many experimental and theoretical efforts (8, 9, 18–20, 10–17). Prior studies found that the nucleosomal DNA is indeed flexible, and unlike the tightly bound configuration observed in the crystal structure, the two ends of the DNA molecule can transiently unwind from the histone core to become exposed on a second timescale (19, 21). Uncontrolled spontaneous unwinding, however, could lead to an irreversible release of the histone core proteins and nucleosome disassembly, jeopardizing the genome stability. Consequently, the nucleosome must tightly regulate its dynamical motion to balance structural stability with DNA accessibility to ensure proper function. The mechanisms which create and maintain this balance, however, are not fully understood.

Single molecule studies have suggested a three-stage model in which the unwinding of the inner and outer layers are decoupled and separated by a large energetic barrier (8, 22). The presence of an energetic barrier will help to keep the inner DNA intact to maintain genome stability as the outer layer unwinds to promote DNA accessibility. However, the lack of structural resolution in single molecule experiments has made mechanistic interpretations of their results non-trivial, and the origin of the barrier that separates outer and inner layer unwinding remains controversial. Often, the crystal structure for the fully wound nucleosome is invoked to highlight the importance of specific chemical interactions for nucleosome stability and the unwinding kinetics (7, 8). However, it is unclear how well observations in this static snapshot generalize to partially unwound nucleosome configurations. Theoretical studies that model the nucleosome as a wormlike chain bound to a cylinder have indeed identified highly twisted DNA conformations near the transition region that differ significantly from the fully wound crystal structure (11, 12). These novel findings have led the authors to argue for the importance of the underlying geometry and physics of the DNA spool in giving rise to the transition barrier. Further first-principles studies are necessary to refine our understanding of nucleosome unwinding by resolving the poorly characterized nucleosome configurations at high resolution.

We applied a chemically accurate, near atomistic protein-DNA model to determine the free energy profile for the entire nucleosome unwinding process as a function of the DNA end-to-end distance. Computer simulations not only quantitatively reproduce thermodynamic parameters determined in the single molecule experiments, but also capture conformational changes of the nucleosome at an atomic resolution as the DNA unwinds. Comprehensive analysis of simulated nucleosome structures suggests that the transition barrier that separates outer and inner DNA unwinding has a strong chemical origin and mainly arises from a delayed loss of histone tails and DNA contacts. Strikingly, this loss of contact mobilizes the disordered tails and leads to a significant increase of entropy that favors nucleosome unwinding. Our study highlights the importance of entropic contributions to nucleosome stability that arise from the conformational flexibility of intrinsically disordered proteins.

## Methods

### Coarse-Grained Protein-DNA Model

In this study we model the nucleosome through the combination of two coarse-grained models: the three sites per nucleotide (3SPN2.C) DNA model and the associative memory, water-mediated, structure and energy model (AWSEM) for proteins (23–25). The 3SPN2.C model is a three-site per nucleotide coarse-grained model for DNA which was parameterized to replicate key mechanical properties such as persistence length, melting temperature, and geometrical properties at varying ionic concentrations. This particular version also accounts for the sequence dependence of these properties through the addition of sequence-dependent terms in the potential (26). In a similar vein, the AWSEM protein model uses C_α_, C_β_, and O sites to represent each residue in a protein (27). Interactions between residues include traditional physically motivated terms such as bonds, angles, dihedrals, etc., as well as bioinformatics terms which bias short (~9-10) sequences of amino acids towards structural motifs commonly observed in other, similar proteins. This model has been successfully employed for structure prediction of monomeric proteins (27), prediction of protein-protein interfaces when monomer structures are known beforehand (28), and investigation of protein misfolding and aggregation (29).

While each of these models contains highly optimized self-interactions, additional force-field terms must be introduced to describe DNA-protein interactions. We introduce non-specific electrostatic interactions at the Debye-Hückel level and Lennard-jones interactions to account for excluded volume effects. Although only a first-order approximation to the true electrostatic environment of the system, such an approach is justified due to the lack of base-specific interactions between the histones and DNA (30). Detailed simulation parameters can be found in the supporting information.

### Simulation Details

We use the end-to-end distance of the nucleosomal DNA as the reaction coordinate to study the thermodynamics of nucleosome unwinding. Formally, we define the ends of the DNA as the center of mass of the sugar sites in the first or last five base pairs and the end-to-end distance as the Euclidean distance between them. This choice of coordinate provides an intuitive metric for the unwinding process and allows for natural comparison with the nucleosome extension coordinate used in single-molecule pulling experiments.

To ensure sufficient exploration of configurational space, we employ umbrella sampling (31) and temperature replica exchange molecular dynamics (32). Fifteen umbrella windows were used, 11 of these were equally spaced from 75 Å to 325 Å in 25 Å increments. Four additional windows were added at 190 Å, 240 Å, 265 Å, and 310 Å to facilitate sampling of specific regions of the reaction coordinate. The spring constant, *k_w_* in kcal/mol·Å^2^ for an umbrella window centered at an end-to-end distance of *d* angstroms was chosen as follows:

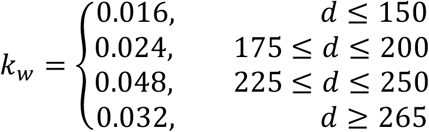

The increase in spring constant at longer distances improves the sampling efficiency at regions with large free energy barriers. Twelve replicas were used within each umbrella window. Temperatures for these replicas ranged from 260 K to 370 K in 10 K increments. Exchanges between these replicas were attempted every 100 timesteps.

All simulations were performed under constant temperature and volume without periodic boundary conditions using the Large-Scale Atomic/Molecular Massively Parallel Simulator (LAMMPS) software package (33, 34). A significant amount of computational effort was exerted to ensure sufficient sampling; almost all umbrella windows were simulated for over 2 × 10^7^ timesteps (Table S1). A coarse-grained representation of the nucleosome constructed from the crystal structure (PDB ID: 1KX5) was used to initialize the simulations. Without losing generality, we used the sequence provided in the crystal structure for the DNA molecule. See the text for a discussion on the robustness of our simulation results with respect to the choice of the DNA sequence.

### Tailless nucleosome system

An additional, independent series of simulations were performed for a tailless nucleosome. Histone tails were removed as described in the experimental study conducted by Brower-Toland et al (9). As with the intact system, umbrella sampling and temperature replica exchange molecular dynamics were used to enhance sampling. For these simulations thirteen umbrella windows were used, 11 of them equally spaced from 75 Å to 325 Å in 25 Å increments. Additional windows were added at 110 Å and 280 Å to ensure the end-to-end distance coordinate was completely sampled. These spring constants, *k_w_*, in kcal/mol·Å^2^ were as follows:

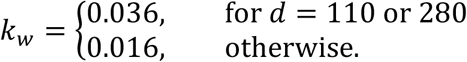

Replica temperatures and attempted exchange rate were identical to those in the intact simulations.

### Thermodynamic Calculations

Unless otherwise specified, free energies and expectations were calculated using the pymbar package developed by the Chodera group (35). Calculations were performed using simulation data from all temperatures subsampled to produce statistically independent samples. Statistical inefficiency was capped at 50 timesteps as high-temperature systems could exhibit long timescale processes such as histone disassembly which were not relevant to the unwinding process.

To calculate expectations as a function of the end-to-end distance, we chose one hundred distance values evenly spaced between 70 and 330 Å and defined a new thermodynamic state for each of them. Each state was assigned a harmonic potential centered at the given distance; the spring constant for this potential was set to an arbitrarily high value (>20 times that used for a typical simulation window potential). Computing expectations of each of these states, therefore, yield thermodynamic expectations at a specific end-to-end distance, thereby allowing us to map out the expected values across the distance spectrum. Potentials of mean force are calculated similarly. The expectation of an indicator function is determined for each bin along the reaction coordinate, and the resulting probability is then used to calculate the free energy. Uncertainties in estimating these quantities can be directly assessed through the calculation of an asymptotic covariance matrix, which is automated by the pymbar package (35).

### Nucleosome Coordinate System

The coordinate system for studying histone tail distribution is defined as follows. First, we locate the DNA intersection point where the two endpoints of the inner layer DNA meet. We note that the inner layer cannot be simply defined based on the crystal structure due to the presence of asymmetric unwinding. Instead, we defined the inner layer as the 81-base pair long DNA segment that contains the most bound base pairs. An 81-base pair region corresponds to approximately one full wrap around the nucleosome. To begin, we determine which of the 147 base pairs are bound using a distance cutoff of σ = 55.0 Å

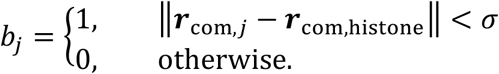

Here **r**_com,j_ indicates the center of mass of the j^th^ base pair and **r**_com,histone_ is the center of mass of the histone octamer. The number of bound base pairs for a continuous stretch of 81-base pair long DNA segment whose first base pair starts as *l* is determined as: 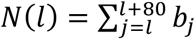. The index for the first base pair of the inner layer *k* is then determined as the *l* with largest *N*(*l*)

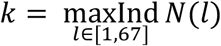

Here, possible values for *l* are constrained by the total length of the nucleosomal DNA that is 147 base pairs long. By definition, *k* + 80 is the index of the last base pair for the inner layer. Note that in the case there are multiple windows with the same number of bound base pairs we chose the average index rounded to the nearest integer. We then define the position of the overlap region being the center of mass of these two base pairs.

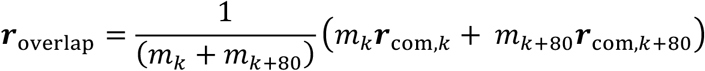

where *m_k_* denotes the mass of the k^th^ base pair.

Next, we calculate the vertical direction of the nucleosome via the average cross product between sequential segments of the inner DNA layer.

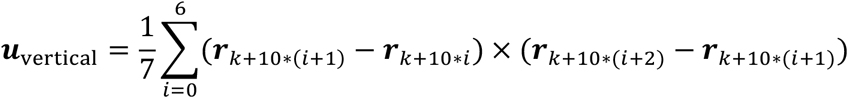

Finally, we can define our coordinate system as follows

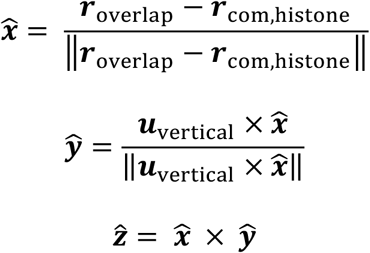

### Data availability

Trajectories and thermodynamic data are available upon request.

## Results

### Free energy landscape of nucleosome unwinding

To provide mechanistic insights into the DNA unwinding process, we mapped out the free energy profile as a function of the end-to-end distance using a coarse-grained protein-DNA model. The end-to-end distance mimics the nucleosome extension manipulated in typical singlemolecule force spectroscopy experiments (10, 36, 37). We chose the coarse-grained model for its chemical accuracy that has been extensively tested in prior studies (27, 38–40) and for its computational efficiency which enables long timescale conformational sampling.

As shown in Figure 1, the free energy increases monotonically as the end-to-end distance stretches from 80 to 320 Å, a range that covers the unwinding of both outer and inner layer of nucleosomal DNA (see Figure 1 insets). Complete dissociation of histone core proteins at distances larger than 320 Å can become irreversible and is not modeled in our simulation. Notably, features along the free energy profile support the three-stage model proposed by Wang and co-workers (8). In particular, the minimum at 80 Å corresponds to fully wound nucleosome conformations which closely resemble the one captured by the crystal structure (7). The plateau region around 200 Å corresponds to partially unwound nucleosome configurations with the DNA outer layer exposed. Finally, the inner layer nucleosomal DNA begins to unwind at distances larger than 250 Å.

**Figure 1.**
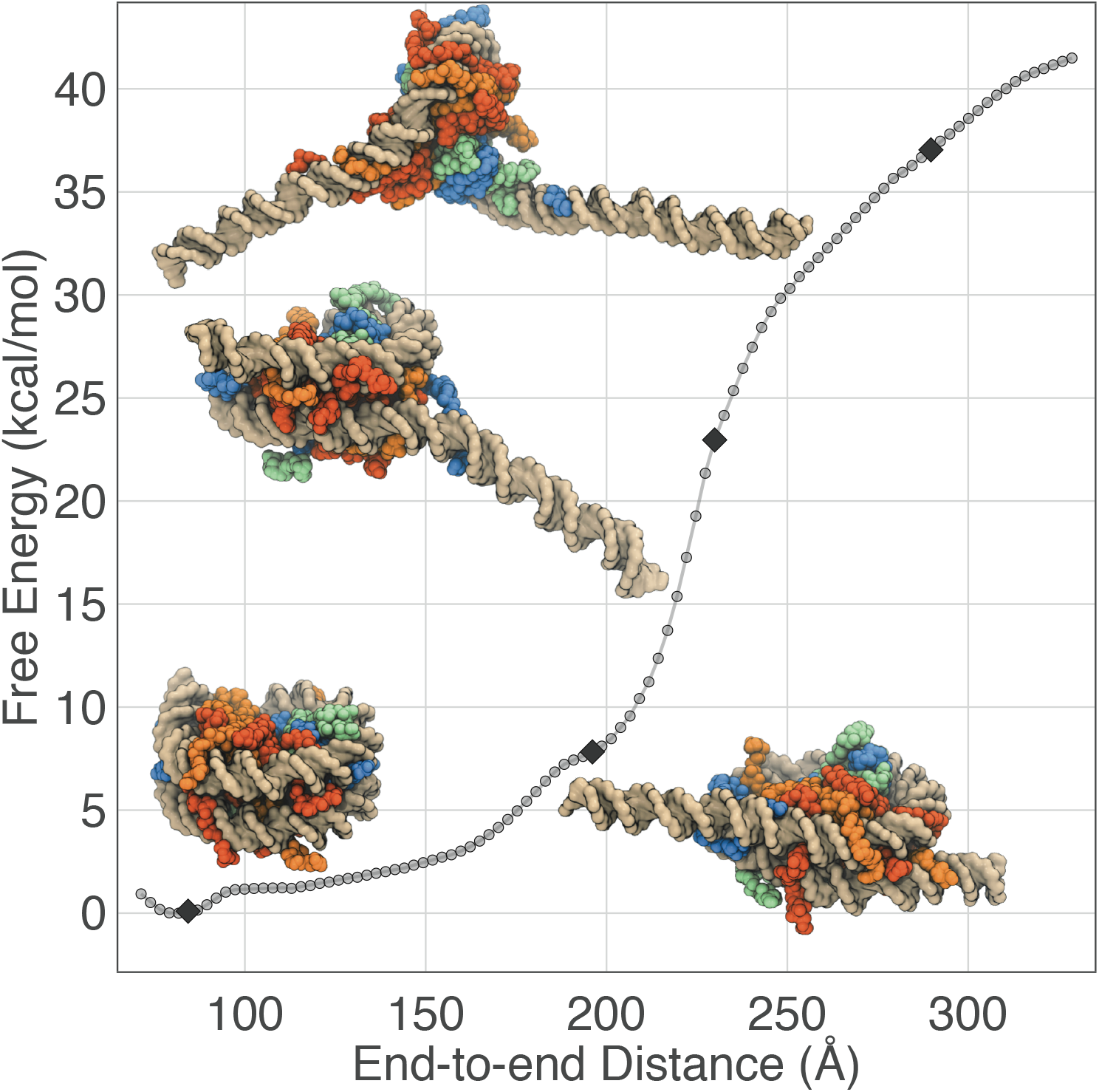
Free energy profile of nucleosome unwinding as a function of the DNA end-to-end distance. Error bars are smaller than symbols. Inset: example nucleosome configurations from end-to-end distances of roughly 80 Å (bottom left), 195 Å (bottom right), 230 Å (center), and 290 Å (top). The DNA molecule in these structures is shown in golden surface, and the histone proteins are drawn in blue (H3), green (H4), orange (H2A), and red (H2B).

Further analyses suggest that the simulated free energy profile quantitatively reproduces energetic barriers between different states as determined in single-molecule pulling experiments. It has been reported that the energetic cost of unwinding the outer layer of nucleosomal DNA is approximately 10 kcal/mol (8, 10, 41), which is in excellent agreement with the free energy difference between 200 Å and 80 Å. Importantly, we observe a sharp increase in free energy as the system transitions from outer to inner layer DNA unwinding in the region from 200 Å to 250 Å, resulting in a total barrier of approximately 20 kcal/mol. The rise of this barrier signals the transition to a high force region that has been detected in numerous single-molecule pulling experiments (8, 10), as can be seen in the gradient of the free energy profile shown in Figure S1. The magnitude of the barrier in this region is also in quantitative agreement with previous experimental and theoretical studies (10, 13, 42). We note that exactly matching simulation and experimental results can be challenging since the large DNA handles used in experiments may impact the barrier for unwrapping.

Close inspection of the simulated structural ensemble suggests that many qualitative observations of the unwinding process are captured by the coarse-grained model as well. Consistent with single-molecule Förster resonance energy transfer (FRET) (15, 16) and cryo-electron microscopy studies (43), we observed large-scale rearrangements of protein-protein interfaces in the histone octamer as the DNA unwinds (Figure S2). In fact, a significant fraction of the histone proteins disassembles into H3-H4 tetramer and two H2A-H2B dimers at end-to-end distances larger than 300 Å (Figure S2). It's worth noting that our protein model correctly captures the stability of the histone octamer at physiological and high salt concentrations (Figure S3). Finally, we confirm the presence of both symmetric and asymmetric unwinding pathways in which either equal amounts of DNA are released from the two ends, or the nucleosome preferentially unwinds one end (17, 38, 44) (see Figure S4). The excellent agreements between modeling and experiments thus justify a further analysis of simulation results to reveal the underlying molecular mechanism of nucleosome unwinding.

We note that the results presented above are a significant generalization of our prior study (38) by extending the DNA unwinding to a much wider range. Both inner and outer layer unwinding can be observed in the end-to-end distances studied here, leading to the discovery of a transition region with a sharp energetic barrier that separates the two.

### Entropic contributions to nucleosome unwinding

We next set out to further characterize the transition region (from 200 to 250 Å) that separates the unwinding of the outer and inner layer of the DNA. The mechanistic origin for the emergence of a large energetic barrier remains controversial, and multiple explanations have been proposed which attribute the barrier to either chemical interactions among histone proteins and the DNA (8, 9) or mechanical deformation of the DNA (12).

We first attempted to understand the shape of this free energy profile by considering the two most intuitive contributions to the energetics of the system - the unwinding of the DNA strand and the electrostatic interactions between the positively charged histone and the negatively charged phosphates of the DNA backbone. The length of the 147 base-pair DNA strand used in this study is on the same order as the persistence length (45). Therefore, nucleosomal DNA is under substantial mechanical stress, and relaxation from the initially highly bent state will decrease the total energy. As this unwinding occurs, the DNA is pulled away from the histone, weakening the favorable electrostatic contacts between them. To test if these two energetic contributions are sufficient to reproduce the free energy profile shown in Figure 1, we calculated the average energy as a function of end-to-end distance for each term as well as their sum; the result is shown in Figure 2.

**Figure 2.**
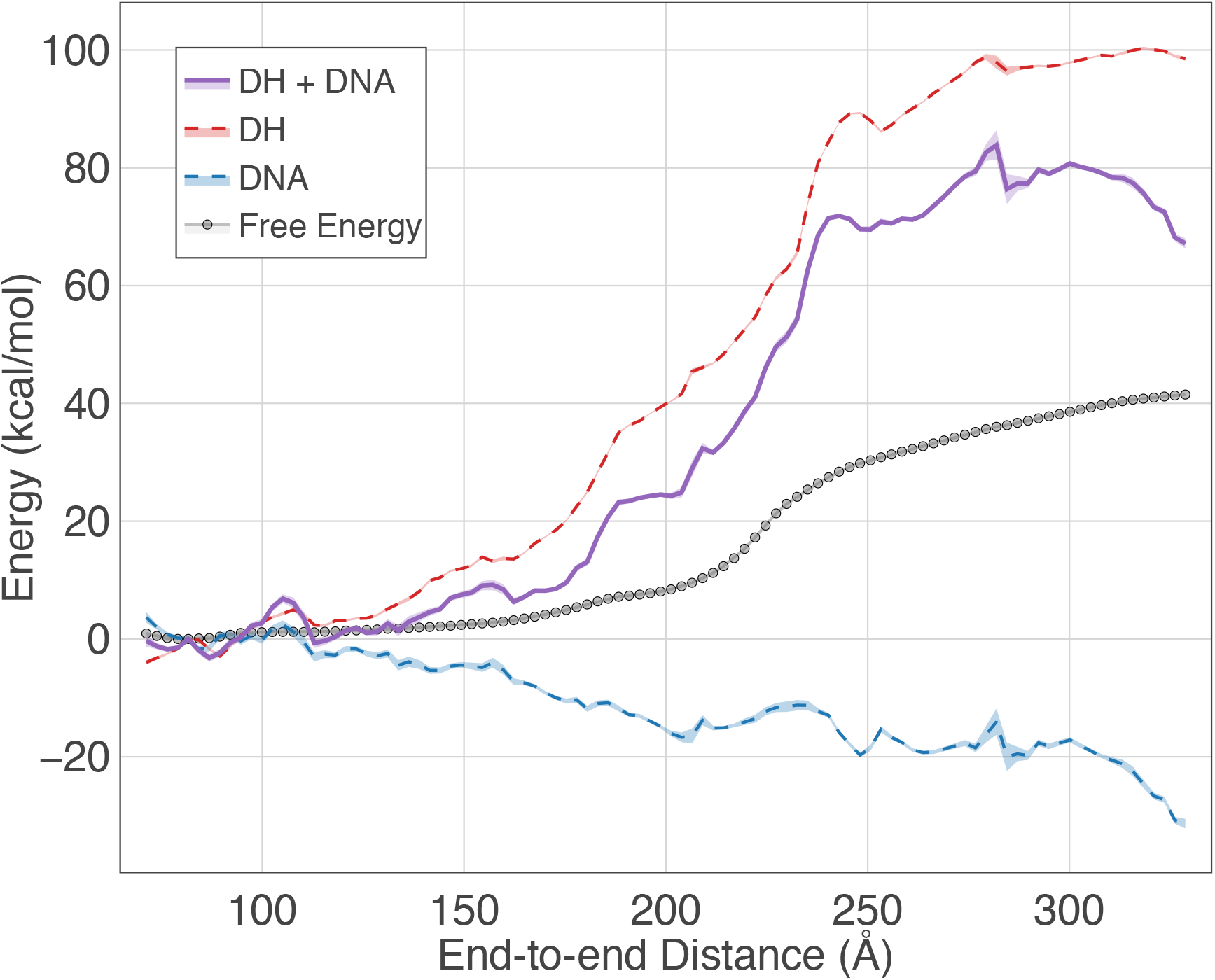
comparison of calculated free energy profile with the sum of average protein-DNA electrostatic (DH) and DNA internal (DNA) energies. Error bars are shown as shaded regions around each curve.

By summing the average energetic contributions from the DNA bending and electrostatics we roughly capture the general shape of the free energy barrier, but dramatically overestimate its magnitude. The barrier in the 200-250 Å region is approximately 40 kcal/mol, double the height of the transition barrier in the free energy profile. Moreover, we fail to accurately capture the free energy change over end-to-end distances 135-200 Å. Consequently, there must be at least one additional aspect of the system that makes a favorable free energy contribution as we increase the end-to-end distance. Presumably, this contribution could be due to any of the other force-field terms that we have neglected to consider thus far, such as the protein energy, nonelectrostatic protein-DNA interactions, etc. Accordingly, we repeat this analysis for all remaining force-field terms and project their average onto the end-to-end distance coordinate. Surprisingly, this does not yield an appreciable change in the energy - the DNA and electrostatic terms alone capture the majority of the total potential energy change (Figure S5). Consequently, we assume there must be a significant entropic contribution to the free energy of unwinding.

Given that the simulations were performed using a temperature replica exchange scheme (32), we can compute the free energy profile at each replica temperature and estimate the relative entropic contribution at 300 K through the finite difference method (46). As illustrated in Figure 3, there is indeed a significant entropic contribution to nucleosome unwinding - the relative entropic component of the free energy decreases by approximately 60 kcal/mol across the end-to-end distance spectrum. If we combine the energetic and entropic components we now recover the calculated free energy profile within error, as shown in Figure 4. Notably, the relative entropic contribution to free energy in the barrier region is approximately -20 kcal/mol, which significantly offsets the energetic penalty (see Figure 4, right panel).

**Figure 3.**
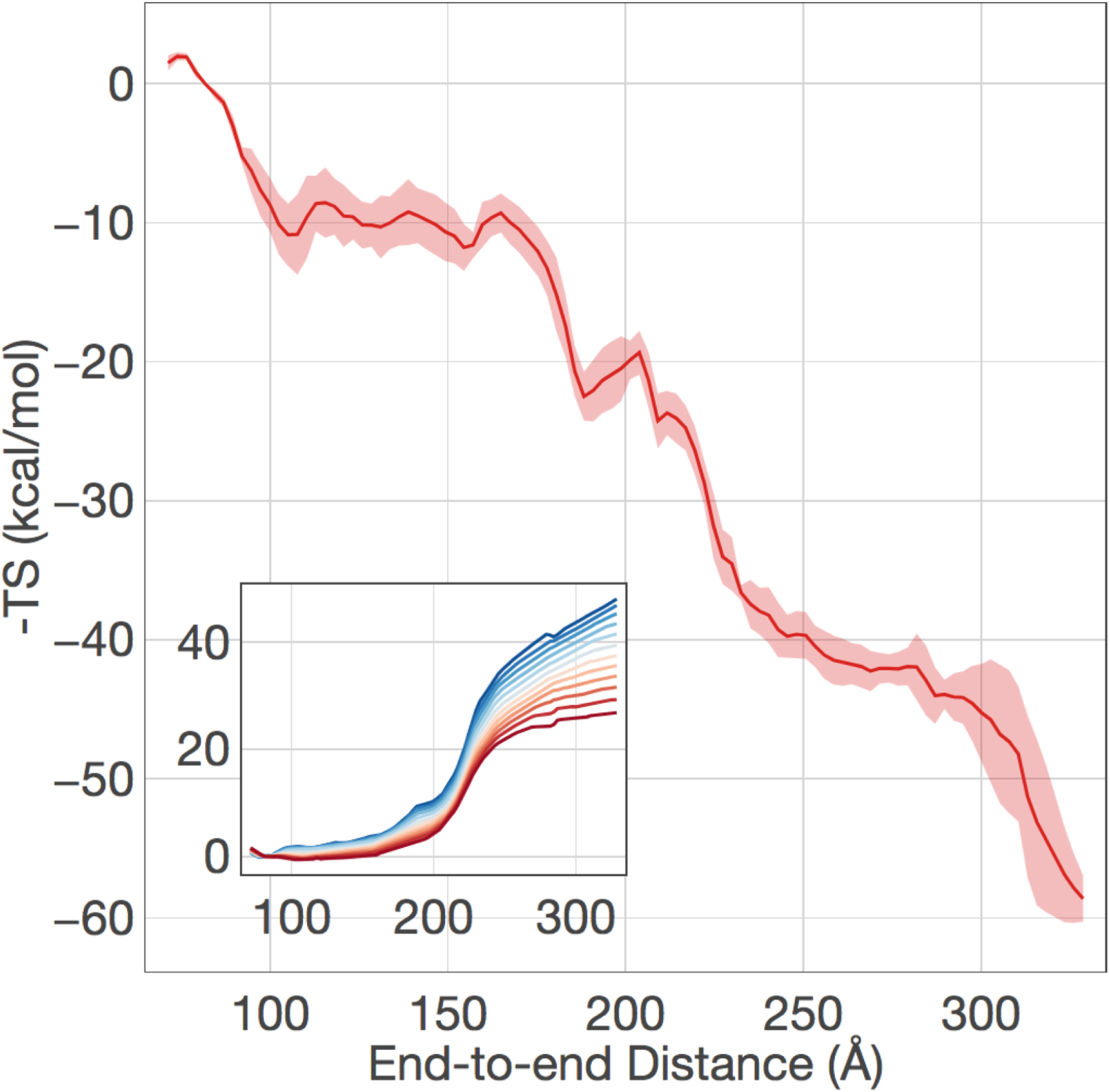
Relative entropic contribution to free energy of unwinding. Entropy was calculated via the finite difference method for all pairs of free energy profiles collected from replicas at 280-320 K. Error calculated as standard deviation of the mean. Inset: free energy profiles calculated from temperatures of 260-370 K. High temperatures are indicated in red, low temperatures in blue.

**Figure 4.**
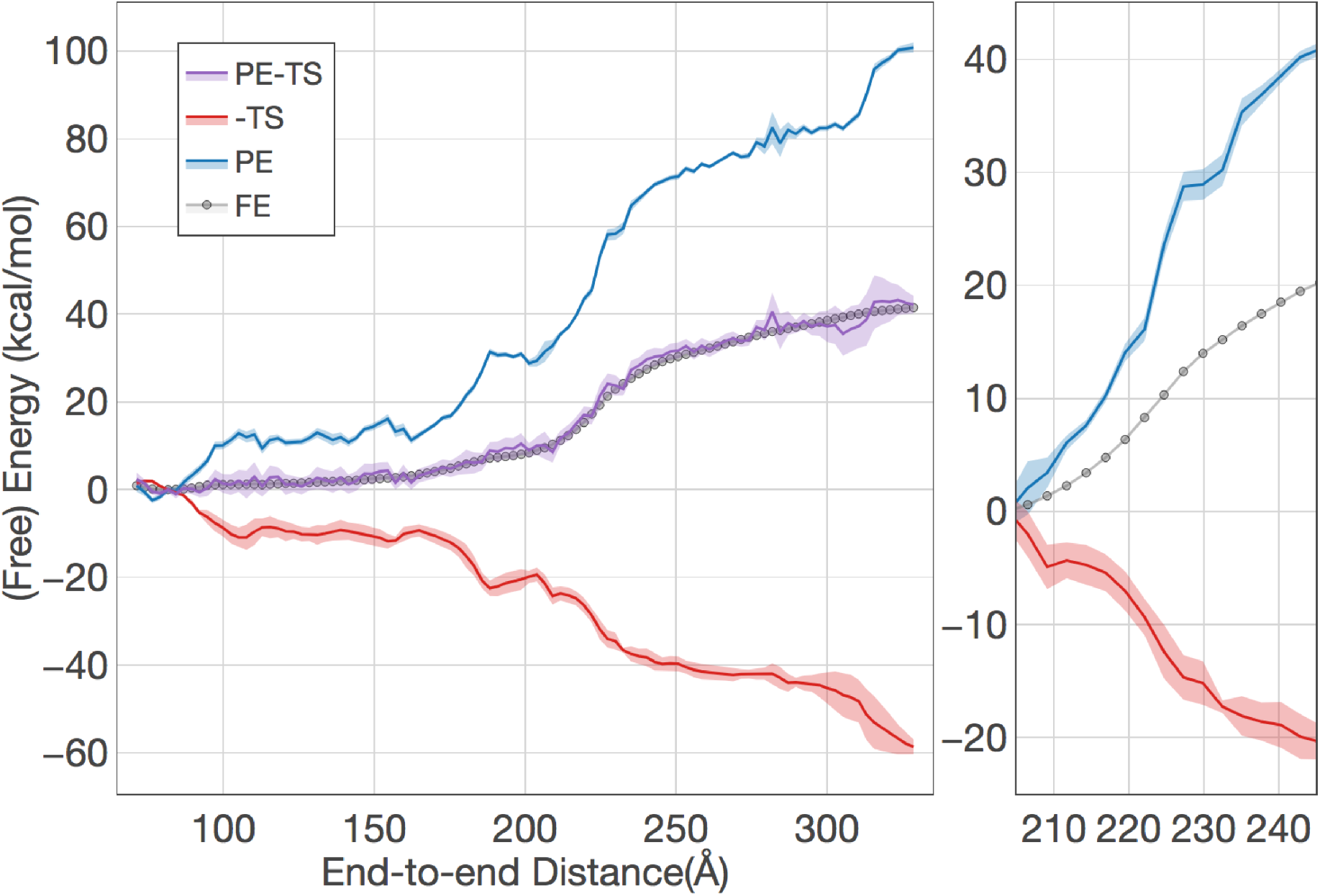
Comparison of free energy (FE) profile with the sum of potential energy (PE) and entropy (-TS) components. The entropy is scaled by the negative temperature (T) for a direct comparison. The sum of potential energy and entropy has been shifted to minimize differences with the free energy profiles as all quantities are only defined up to a constant. Right: an expanded view of the barrier region. All quantities have been shifted to zero at the beginning of the barrier region. The sum of the potential energy and the entropy has been removed for clarity.

We conclude that it is electrostatic interactions that give rise to the energetic barrier in the transition region. This increase in electrostatic energy, however, is offset by a concomitant increase in entropy, resulting in an overall modest change in terms of free energy. We note that these conclusions should be robust with respect to the particular DNA sequence used in the simulation. We used the sequence provided in the PDB structure, but variations in the sequence will give rise to a change in the DNA bending energy that is at most of 6 kcal/mol (47). This variation is much smaller than the energetic and entropic contributions in the transition region that are over 20 kcal/mol.

### Histone tails safeguard the inner layer

To understand the origin of the surprisingly large entropic contribution to the transition region, we further characterized structural changes of the nucleosome as the DNA unwinds. We note that near the transition region, the length of the unwound DNA is approximately 70bp (see Figure S6), which is well below the persistence length. Furthermore, the histone core protein remains stable in conformations that closely resemble the crystal structure (see Figure S2). The contribution from these two components to the entropy is thus expected to be small, and we, therefore, focused our analysis on histone tails.

To directly investigate the mobility of histone tails, we defined a structure-specific coordinate system where the x-y plane coincides with the nucleosomal plane (Figure 5A). The x-axis points from the center of the nucleosome to the DNA intersection, which is defined as the location where the two endpoints of the inner layer DNA meet (see *Methods: Nucleosome coordinate system for details*). In Figure 5B-5D, we plot the probability density distributions of the position for the last residue of histone tails onto the x-y plane at different end-to-end distances. At the beginning of transition region where the end-to-end distance is approximately 200 Å, histone tails strongly localize at the intersection region (x ~ 25 Å) as indicated by the high probability density. Near the end of the transition region, however, histone tails delocalize over the entire nucleosomal plane, and the probability distribution is more uniform. The significant variation in their conformational flexibility thus supports the role of histone tails in contributing to the entropy rise along the transition region.

**Figure 5.**
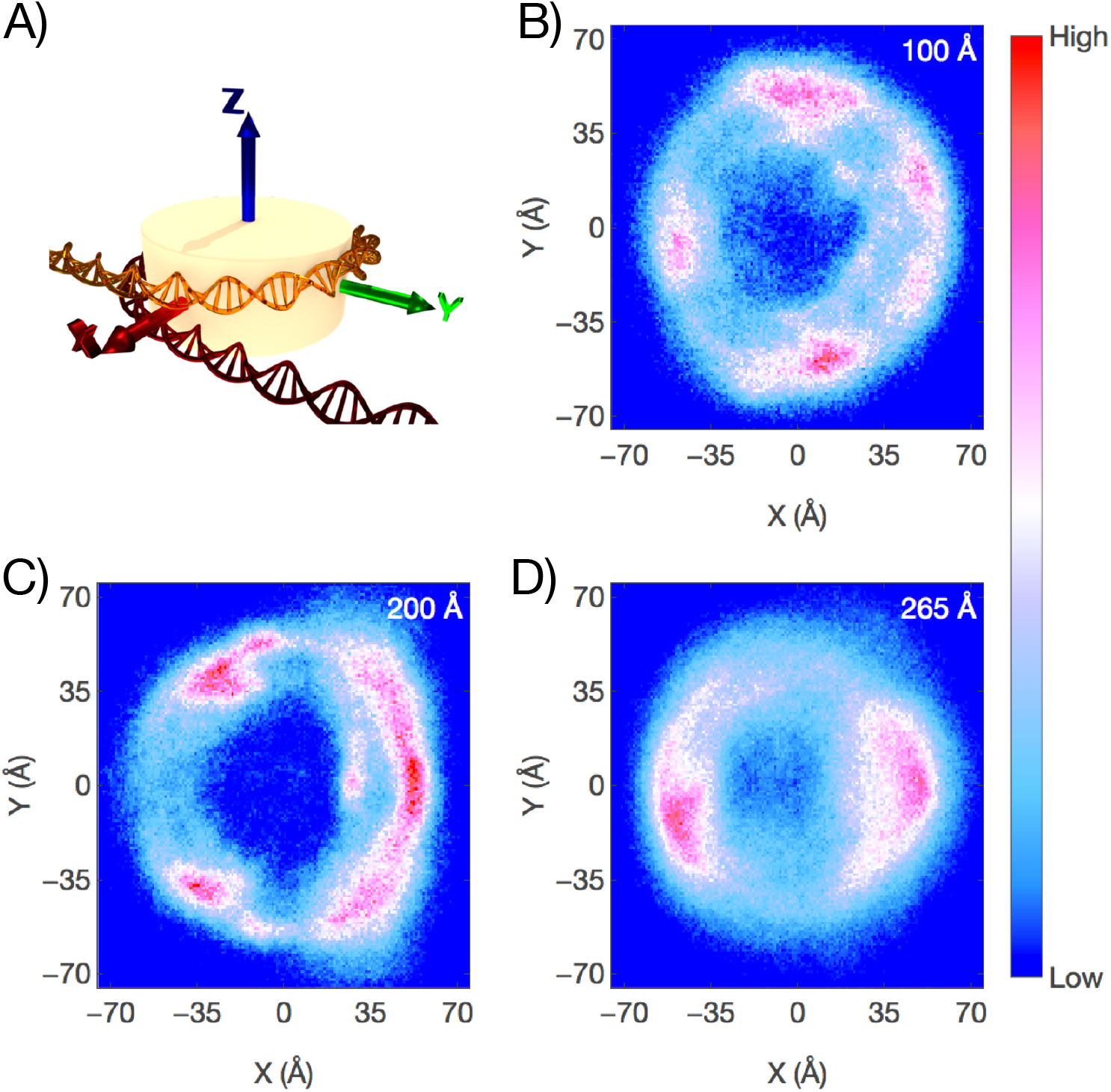
Histone tail position distributions relative to the DNA overlap region. (A) Schematic representation of the nucleosome coordinate system (see Methods section for additional details). (B-D) Projections of the last alpha carbon in each histone tail onto the x-y plane for end-to-end distances 100 Å (top right), 200 Å (bottom left), and 265 Å (bottom right). Each window contains data from all trajectory frames with end-to-end distances within ±5 Å of the listed value.

A substantial entropic contribution from histone tails would be consistent with the change of their interaction energy with the DNA as a function of the end-to-end distance. As shown in Figure S7, similar to the total electrostatic energy, the histone tail-DNA energy increases sharply in the transition region, and in fact, accounts for most of the changes observed in Figure 2. On the contrary, a more gradual change for the histone core-DNA energy is observed in the same region. Further decomposing the tail-DNA energy into individual proteins suggests that the energy comes from interactions between H2A and H2B with the DNA, though considerable contributions from H3 and H4 are also observed (see Figure S7). Breaking the tail-DNA contacts will significantly mobilize histone tails, giving rise to a large increase in entropy in the transition region.

The above analysis indicates that it is the sudden loss of tail-DNA contacts that gives rise to the large jump in both energy and entropy observed in the transition region. Remarkably, histone tails manage to preserve their interaction with the DNA molecule until the two DNA ends reach a separation of 200 Å. This delay in tail contact loss is a result of large-scale rearrangements of histone tails that are most evident by comparing the probability density distributions at 100 and 200 Å. Multiple peaks can be seen in the distribution at 100 Å (Figure 5B), indicating the strong binding of histone tails with different regions of the nucleosomal DNA. On the other hand, most of the histone tails localize at the intersection region as the end-to-end distance reaches 200 Å. The drive for this dynamical reallocation of histone tails is to search for nucleosomal regions with two strands of DNA to maximize favorable electrostatic interactions. This rearrangement helps to keep both the energy and entropy relatively constant until the transition point, the place at which the last region with two strands of DNA is lost. This rearrangement, of course, cannot perfectly preserve all contact energies, and the entropic increase before 100 Å indeed arises from a partial loss of contact between histone H3 tails and the DNA (see Figure S7).

Consistent with the above interpretation that emphasizes the central role of histone tails in safeguarding the inner layer from unwinding, we find that for a tailless nucleosome, thermodynamic features corresponding to the transition region become less prominent. The tailless nucleosome is designed by removing segments of histone tails to mimic the product of trypsin digestion (9). We carried out independent simulations and analysis to characterize its free energy profile as a function of the end-to-end distance. As shown in Figure 6 and Figure S1, the variation in the slope of the free energy profile for end-to-end distances larger than 200 Å is significantly reduced relative to the intact system. Furthermore, the energetic cost from 200 to 250 Å is about 15 kcal/mol, reduced considerably from the 40 kcal/mol seen in the intact system. Similarly, the relative entropy variation in this regime is barely noticeable.

**Figure 6.**
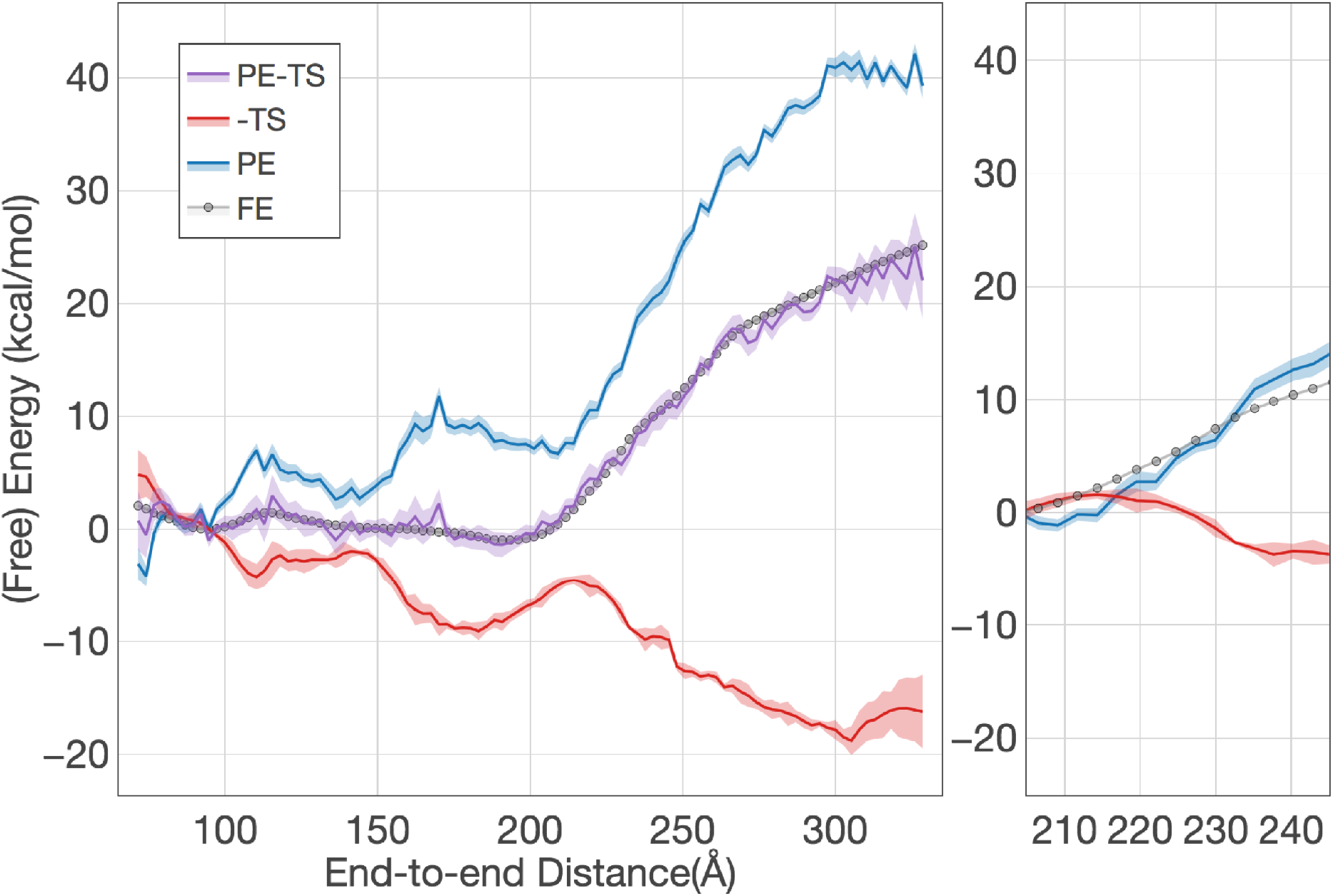
Comparison of free energy (FE) profile with the sum of potential energy (PE) and entropy (-TS) components for the tailless system. The entropy is scaled by the negative temperature (T) for a direct comparison. Right: an expanded view of the barrier region. All quantities have been shifted to zero at the beginning of the barrier region. The sum of the potential energy and the entropy has been removed for clarity.

### Histone tails regulate asymmetric outer layer unwinding

In addition to their major contributions to the transition region, histone tails play a critical role in the unwinding behavior of the outer layer of the DNA. Specifically, consistent with previous experimental and computational studies (17, 38), we observed a predominantly asymmetric pathway for the unwinding of nucleosomal DNA. This asymmetry is most evident from the twodimensional free energy profiles provided in Figure S4. The two axes measure the change in the radius of gyration of the first (base pairs 1 to 73) and second (base pairs 74 to 147) half of the DNA molecule respectively. An increase in the radius of gyration is expected if the corresponding DNA segment unbinds from the histone core. Along asymmetric pathways, the two DNA segments are relatively uncoupled, allowing the nucleosome to preferentially expose one DNA end while keeping the other one fixed. This is in contrast to the symmetric pathway in which the motions are more concerted and both DNA ends are exposed at the same rate. Note that since the DNA sequence studied here is palindromic, the two asymmetric pathways have identical thermodynamic weight, as can be seen from the symmetry of the free energy profiles. Illustrative nucleosome configurations along different unwinding pathways are provided in Figure 7A.

**Figure 7.**
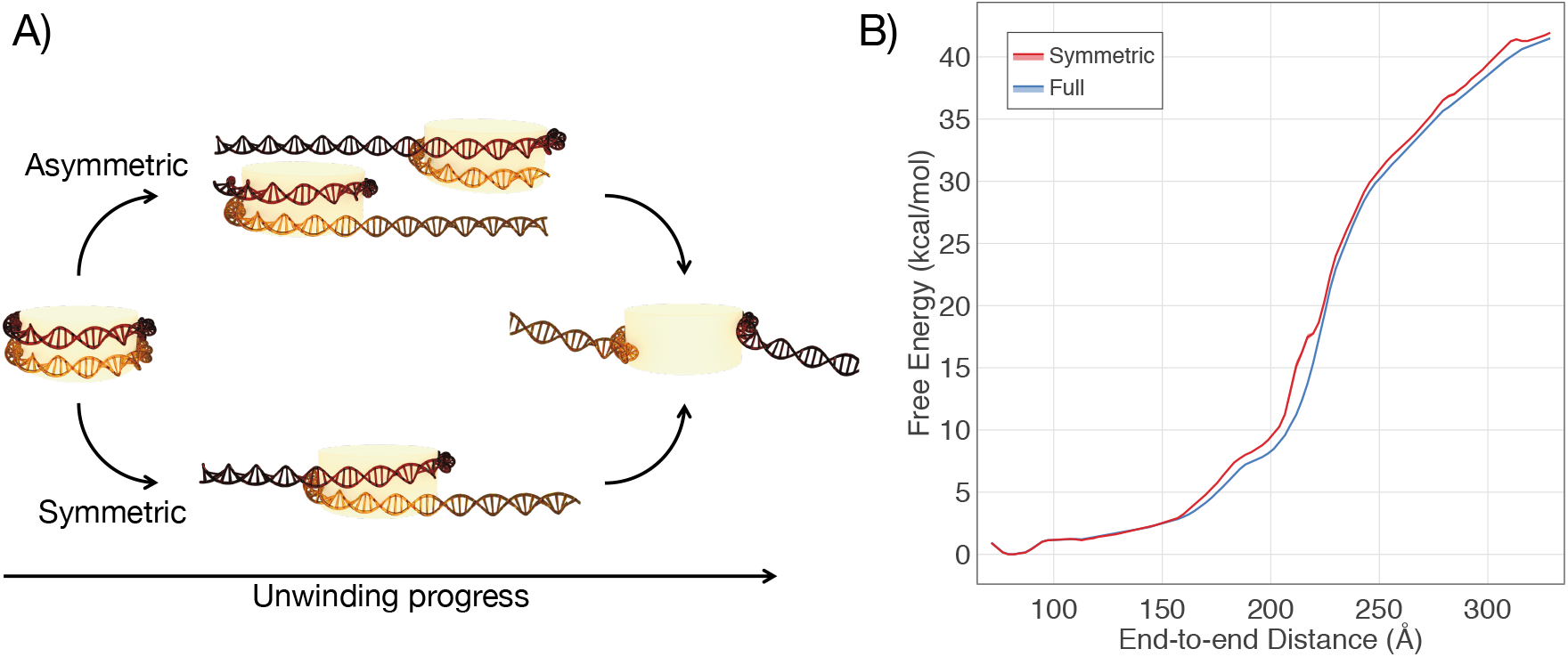
Illustration and comparison of the asymmetric and symmetric pathways for nucleosome unwinding. A) Schematic diagram of the two pathways observed during unwinding. B) Comparison of the free energy profile shown previously with the free energy of unwinding in an exclusively symmetric fashion.

Though both pathways are present, the DNA unwinds predominately in the asymmetric fashion for the intact system. To better understand the thermodynamic stability of different pathways, we computed the free energy for unwinding exclusively in a symmetric fashion and compared this to the full free energy (see Figure 7B). Symmetric unwinding of an intact nucleosome resulted in a free energy 1 to 2 kcal/mol greater than that of the full free energy profile, therefore explaining the strong preference for asymmetric unwinding. We defined symmetric unwinding as nucleosome configurations in which the difference between the radius of gyration for the two DNA halves is less than 10 Å. Strikingly, we find that the preference of asymmetric pathway is dependent on the presence of histone tails, and is essentially non-existent for the tailless system. As shown in Figure S8, the free energy difference between the symmetric and total free energy profiles to be significantly smaller (<0.5 kcal/mol) once histone tails are removed.

We further decomposed the free energy difference shown in Figure 7B into energetic and entropic contributions for additional mechanistic insights (see Figure S9). The symmetric unwinding is energetically more unfavorable for almost all end-to-end distances, though the energetic penalty is partially offset by entropic contributions. Further analysis suggests that the increased energetic cost arises mostly from the presence of more protein deformation along the symmetric pathway (see Figure S10C and S10D). Previously, the sequence dependent DNA mechanics has been invoked to rationalize the relative stability of the two asymmetric configurations (17, 44). We argue here that for the preference of asymmetric configurations over symmetric ones, other factors in addition to DNA mechanics, such as octamer stability, play a role as well. In fact, as shown in Figure S10B, the DNA energy is actually lower in symmetric configurations, even though the symmetric pathway is overall unfavorable.

## DISCUSSION

We have shown that a coarse-grained protein-DNA model quantitatively reproduces the free energy profile of single nucleosome winding, validating it for a more detailed investigation of the unwinding mechanism. We find that considering only the energetic contributions from the DNA and electrostatic interactions between the DNA and histone core qualitatively reproduce the shape of the free energy curve, but fails to correctly capture the magnitude of the barrier in the transition region that separates the unwinding of the inner and outer layer of the nucleosomal DNA. Comparing free energy profiles computed at different temperatures indicated there is a significant entropic contribution to the free energy, most notably within the transition region. The molecular origin of this entropy is the enhanced mobility of the flexible histone tails following their unbinding. The free energy cost of nucleosomal DNA unwinding, therefore, is primarily due to electrostatic interactions which are significantly offset by a concomitant increase in entropy generated by breaking these contacts. We validate this explanation through the introduction of an additional set of simulations in which the histone tails have been removed. Overall, both the energetic cost and entropic compensation for DNA unwinding are significantly reduced, leading to a milder free energy profile as the DNA end-to-end distance increase.

The importance of histone tail entropy highlighted in this study provides a fresh view on the role of histone modifications in regulating nucleosomal dynamics. Post-translational modifications of histone tails are widespread and are known to greatly impact the stability of nucleosome structure (9, 38, 48–51). For example, acetylation can neutralize positive charges on these tails and weaken their interaction with the DNA molecule (52). Methylation, on the other hand, has been shown to facilitate the binding of additional chromatin regulators that can drive conformational changes in the nucleosome by utilizing chemical energy (53). In addition to these mechanisms, however, histone modifications could regulate nucleosome stability by fine-tuning the entropic contributions of disordered tails to the free energy cost of DNA unwinding. Indeed, numerous studies have shown that post-translational modifications greatly affect the conformational flexibility of histone tails (54, 55). Further quantitative investigations on how histone tail entropy is affected by these modifications will enhance our understanding of their role in regulating nucleosome structure and function.

## ACKNOWLEDGMENTS

This work was supported by startup funds from the Department of Chemistry at the Massachusetts Institute of Technology. T.P. acknowledges the Stephen J. Lippard Fellowship for financial support.

## Author Contributions

T.P. and B.Z. performed research, analyzed data and wrote the manuscript.

